# Climate shapes flowering periods across plant communities

**DOI:** 10.1101/2021.10.10.463841

**Authors:** Ruby E. Stephens, Hervé Sauquet, Greg R. Guerin, Mingkai Jiang, Daniel Falster, Rachael V. Gallagher

**Affiliations:** Department of Biological Sciences, Macquarie University, Sydney, Australia; National Herbarium of New South Wales (NSW), Royal Botanic Gardens and Domain Trust, Sydney, Australia; Evolution and Ecology Research Centre, School of Biological, Earth and Environmental Sciences, University of New South Wales, Sydney, Australia; Terrestrial Ecosystem Research Network, School of Biological Sciences, The University of Adelaide, Adelaide, Australia; Hawkesbury Institute for the Environment, Western Sydney University, Sydney, Australia

**Keywords:** community assembly, climate, floral traits, flowering phenology, functional biogeography, macroecology, predictability

## Abstract

**Aim:** Climate shapes the composition and function of plant communities globally, but it remains unclear how this influence extends to floral traits. Flowering phenology, or the time period in which a species flowers, has well-studied relationships with climatic signals at the species level but has rarely been explored at a cross-community and continental scale. Here, we characterise the distribution of flowering periods (months of flowering) across continental plant communities encompassing six biomes, and determine the influence of climate on community flowering period lengths.

**Location:** Australia

**Taxon:** Flowering plants

**Methods:** We combined plant composition and abundance data from 629 standardised floristic surveys (AusPlots) with data on flowering period from the AusTraits database and additional primary literature for 2,983 species. We assessed abundance-weighted community mean flowering periods across biomes and tested their relationship with climatic annual means and the predictability of climate conditions using regression models.

**Results:** Combined, temperature and precipitation (annual mean and predictability) explain 29% of variation in continental community flowering period. Plant communities with higher mean temperatures and lower mean precipitation have longer mean flowering periods. Moreover, plant communities in climates with predictable temperatures and, to a lesser extent, predictable precipitation have shorter mean flowering periods. Flowering period varies by biome, being longest in deserts and shortest in alpine and montane communities. For instance, desert communities experience low and unpredictable precipitation and high, unpredictable temperatures and have longer mean flowering periods, with desert species typically flowering at any time of year in response to rain.

**Main conclusions:** Our findings demonstrate the role of current climate conditions in shaping flowering periods across biomes, with implications under climate change. Shifts in flowering periods across climatic gradients reflect changes in plant strategies, affecting patterns of plant growth and reproduction as well as the availability of floral resources across the landscape.

## 1 INTRODUCTION

Climate shapes patterns of community assembly globally, driving the distribution of resources and the dynamics of interactions that in turn affect the co-occurrence of organisms (Kraft et al., 2015; Ockendon et al., 2014). As community composition varies along environmental gradients, so do the functional traits of constituent species (Bruelheide et al., 2018; Cornwell & Ackerly, 2009; Wieczynski et al., 2019). For example, plant communities are generally taller in the tropics, and in areas with higher precipitation (Moles et al., 2009), with leaves on average larger in environments which are warm and wet (Wright et al., 2017). Yet less is known about how the traits of flowers vary with climate across biomes, continents or globally.

Previous studies of plant functional biogeography have primarily focussed on a few key traits thought to be central to plant strategies, particularly leaf size and specific leaf area, plant height and seed mass (Andrew et al., 2021; Lamanna et al., 2014; Swenson et al., 2012). While such studies have been extremely productive in describing plant ecological strategies across a wide range of environmental conditions, recent attention has been drawn to the overlooked role that flowers and floral traits play in modulating species interactions and shaping patterns of community assembly (E-Vojtkó, de Bello, Durka, Kühn, & Götzenberger, 2020; Roddy et al., 2020). Despite some evidence suggesting that floral traits may have weaker links to macroclimate and landscape patterns than vegetative traits in general (e.g. Kuppler et al., 2020), flowers and floral traits do respond to biotic and abiotic conditions and thus bear investigation as “response” traits (Caruso, Eisen, Martin, & Sletvold, 2019; E-Vojtkó et al., 2020; S. Lavorel & Garnier, 2002). At the same time floral traits play important roles in ecological communities, mediating sexual reproduction by cross-pollination in flowering plant species and the provision of food and shelter resources for fauna (Fornoff et al., 2017; Lázaro, Gómez-Martínez, Alomar, González-Estévez, & Traveset, 2020). Thus floral traits also bear investigation as “effect” traits for their influence on other trophic levels and ecosystem functions (E-Vojtkó et al., 2020; Sandra Lavorel et al., 2013).

Flower phenology has strong connections to climatic signals, at the individual, population, species and community level (Craine, Wolkovich, & Towne, 2012; Diez et al., 2012; Primack, 1985), and is thus a prime candidate trait for studies of floral functional biogeography. Flowering phenology is a highly labile trait, with a large amount of intraspecific variation between populations experiencing different climatic and biotic conditions (Franks, Sim, & Weis, 2007; Yan, Wang, Chan, & Mitchell-Olds, 2021). Indeed, flowering phenology shifts have been observed in numerous species worldwide in response to climate warming (e.g. CaraDonna *et al*., 2014; Prevéy *et al*., 2019). Flowering phenology also shifts with community composition, and composition-derived variation in flowering time can explain a significant proportion of community flowering periods (though less than intraspecific variation; Park, 2014).

Recent work suggests that interspecific variation in flowering phenology can be detected at a landscape scale. For example, flowering and fruiting periods of Chinese angiosperms with overlapping geographic ranges vary with latitude, elevation and several climatic variables (Du et al., 2020). However, assessments of variation at grid-cell rather than local patch scale can over-estimate the influence of macro-environment on trait signals among co-existing species (Bruelheide et al., 2018). Species with overlapping broad geographic ranges do not necessarily co-occur in communities at a scale where they are likely to interact, and patterns of trait variation may differ significantly when species abundances within ecosystems are taken into account (Wieczynski et al., 2019). It thus remains unclear whether relationships between community flowering phenology and climatic signals apply to community sorting at the local scale.

Here, we characterise the continental distribution of flowering periods in plant communities, and determine the influence of climate on community flowering period lengths. We define flowering period length as the number of months in which each species has been recorded flowering, which is not necessarily equivalent to the flowering durations of populations or individuals. We combine fine-scale plant community richness and abundance data from a network of vegetation plots across Australia (TERN AusPlots (TERN, 2018)) with flowering period data from the AusTraits database (Falster et al., 2021), species descriptions and herbarium records. The Australian continent, though generally low in soil fertility, encompasses a wide array of climatic regimes from cool temperate to tropical. Vast low relief deserts of the arid interior juxtapose areas of higher elevation such as the Great Dividing Range of eastern Australia and higher rainfall habitats with more predictable climates along coastal fringes (Figure 1). Australia has a latitudinal range of >30°accompanied by a strong gradient in mean annual temperatures.

**Figure 1.**
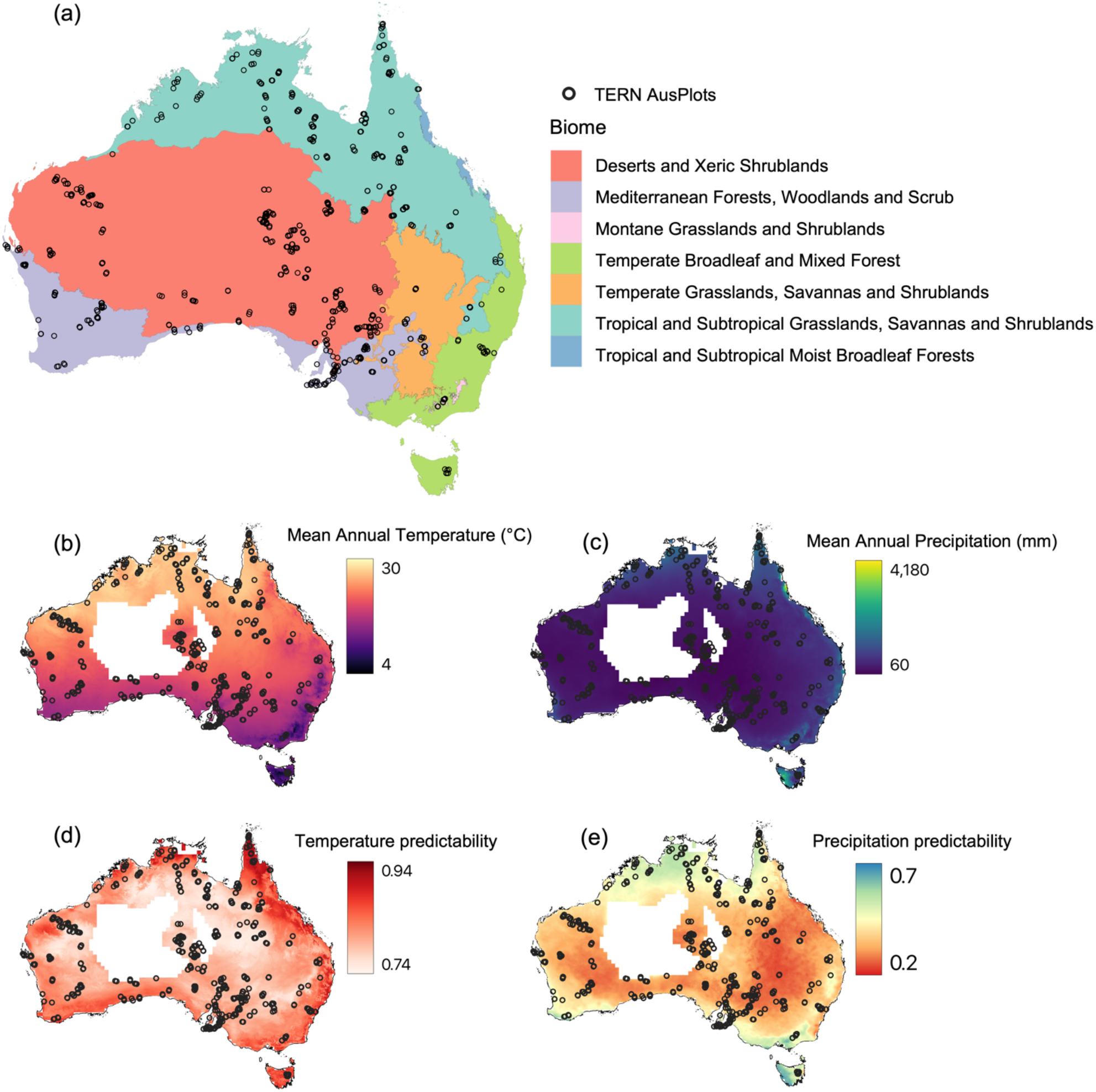
The distribution of the 629 AusPlots used in the analysis, across: (a) biomes based on Dinerstein et al. 2017’s global terrestrial ecoregions, aligned to the Australian Interim Biogeographic Regionalisation for Australia (Australian Department of the Environment and Energy, 2016); (b) mean annual temperature (°C); (c) mean annual precipitation (mm); (d) temperature predictability; and (e) precipitation predictability. Climate data generated from Australian Water Availability Project (AWAP) data for 1930-2019. The white area in central Australia in represents a mask where AWAP data were excluded as meteorological stations are sparse in this area (King et al., 2014).

Climatic conditions may influence the length of community flowering periods in several ways. Higher mean annual temperatures allow pollinators to be active and plants to meet the physiological costs of producing flowers across a longer period of the year (Primack & Inouye, 1993; Roddy, 2019; Roddy et al., 2020), thus lengthening flowering periods. Low mean annual precipitation, on the other hand, reduces water availability and plant productivity which may select for ephemeral flowering strategies (Friedel, Nelson, Sparrow, Kinloch, & Maconochie, 1993; Roddy, 2019), thus also lengthening potential flowering periods overall.

In addition to average climate conditions, we hypothesise that the predictability of climatic phenomena has a strong influence on flowering phenology. We test this idea using the Colwell index of predictability (Colwell, 1974), which combines both the long-term reliability of seasonality, known as contingency, and the constancy of aseasonal periodic phenomena into a single measure of environmental predictability. Predictable environments offer reliable environmental information to organisms, allowing the timing of events such as flowering to depend on endogenous factors such as age or condition rather than responding directly to environmental cues (Wingfield, Hahn, & Doak, 1993). Predictability is therefore a more complete measure of environmental stochasticity than the more commonly used temperature or precipitation seasonality, especially in relatively aseasonal continents such as Australia (Jiang, Felzer, Nielsen, & Medlyn, 2017). Globally temperature predictability follows a latitudinal gradient, and is uniquely high in Australia, with greater predictability closer to the equator and coastal areas (Jiang et al., 2017). Precipitation predictability is more geographically variable, and in Australia is low overall but markedly lowest in the arid inland (Jiang et al., 2017). We expect that high climatic predictability offers reliable environmental cues and therefore selects for synchronous biotic responses, with more concentrated and thus shorter community flowering periods in areas of high predictability. Given that temporal information is preserved by both flowering periods and climatic predictability, we also anticipate that climatic predictability will have a stronger relationship with community flowering period lengths than climatic means.

In summary we predict:

1. That community flowering periods will be longer with increasing mean annual temperature.
2. That community flowering periods will be longer with decreasing mean annual precipitation.
3. That community flowering periods will be shorter with increasing predictability of either temperature or precipitation.
4. That community flowering period length will have a stronger relationship with the predictability of climatic variables than mean climatic measures.

## 2 METHODS

### 2.1 Community floristic data

We accessed data on floristic composition in 810 surveys of 100 m × 100 m vegetation plots from the Terrestrial Ecosystem Research Network (TERN) AusPlots network using the ausplotsR package (Guerin, Munroe, Saleeba, & Ire, 2020; Munroe et al., 2021; TERN, 2021). AusPlots are distributed across a representative range of Australian ecosystems and environments and were surveyed using precise and consistent methods for recording vegetation species and cover-abundance data between 2011-2020 (Guerin, Williams, Leitch, Lowe, & Sparrow, 2021; Guerin, Williams, Sparrow, & Lowe, 2020; Sparrow et al., 2020). Plots were included in analyses if they were located ≥ 500 metres from another plot and flowering period data was available for ≥ 80% of angiosperm species cover (Borgy et al., 2017; Figure S1). Where plots had repeat surveys available, the survey with the highest recorded species richness was retained to maximise representation of species occurring in the system. In total, 629 plots with 2,983 species were retained for analysis (Figure 1). These plots cover a broad and representative range of Australia’s climatic variation (Figure 1) and occur across six globally recognised biomes (Dinerstein et al., 2017, Figure 1). The number of plots sampled in each biome strongly correlates with biome size in Australia (Figure S2, ordinary least squares linear regression *p* < 0.001, R^2^ = 0.92). All observations were aggregated to the species level, removing any subspecies or variants, after taxonomic alignment to the Australian Plant Census (Council of Heads of Australasian Herbaria, 2021) following methods in (Falster et al., 2021)).

### 2.2 Flowering period

Data on flowering periods were accessed from the AusTraits database version 2.1.0 (Falster et al., 2021) drawn from diverse original sources. The data from AusTraits were supplemented for 627 species from species descriptions in the Flora of Australia (Australian Biological Resources Study, Canberra, 2021), online state and regional floras (‘EUCLID’, 2020; Northern Territory Government, 2021; Royal Botanic Gardens and Domain Trust, Sydney, 2021; Royal Botanic Gardens Victoria, 2021; State Herbarium of South Australia, 2021; Western Australian Herbarium, 2021; Zich, Hyland, Whiffin, & Kerrigan, 2020), original species descriptions and, where flowering period was not available from any of the above sources, herbarium records. Most original sources define flowering periods using a range of months, e.g. “Jun-Oct”, “spring-summer” or “all year round”. For analysis, each record was converted into binary vector of length 12, indicating whether flowering occurred in each month, e.g. “110000000011” for Nov-Feb.

Flowering period length was defined as the number of months (i.e. 1-12) in which the species has been recorded flowering. It therefore refers to the proportion of the year during which a species potentially flowers, rather than to the length of flowering events. We use the length of flowering periods as our response variable so as to include the numerous Australian arid-zone species which flower sporadically in response to rain (Friedel et al., 1993; Friedel, Nelson, Sparrow, Kinloch, & Maconochie, 1994). Mean flowering month cannot be calculated for these species as midpoint circular means cannot sensibly be calculated for bimodal or equally spaced periods (Morellato, Alberti, & Hudson, 2010). Where multiple records of flowering period existed for a single species, data were pooled (e.g. a species reported as flowering in both March-April and April-May was scored as flowering March-May). This ensured we captured the full scope of months a species has been reported to flower across its Australian range.

### 2.3 Climatic variables

Climatic variables were calculated for plot locations using CSIRO Australian Water Availability Project (AWAP) data from 1930-2019 (Jones, Wang, & Fawcett, 2009; Raupach et al., 2009, 2012). AWAP temperature and precipitation data use records from the Australian Bureau of Meteorology’s network of meteorological stations across Australia, and are modelled at a resolution of 0.05 degree (~5 km). AWAP data accuracy is reduced for assessments of temporal variability where the meteorological station network is sparse or has missing data, in years before 1930 and in areas in central western Australia and locations along the Australian coast (King et al., 2014). A mask was applied to exclude AWAP data from locations where the network is sparse (as per King et al., 2014; white areas in Figure 1). Fifty-two AusPlots occurred in masked areas and so were excluded from analyses with climatic variables.

Mean annual temperature (°C) (MAT), mean annual precipitation (mm) (MAP) and the Colwell index of predictability (Colwell, 1974) for temperature and precipitation were calculated from AWAP data for each plot location. The Colwell index of predictability is a simple but elegant mathematical approach that condenses temporal patterns of variability into single scores that vary between 0 (completely unpredictable) to 1 (completely predictable). The index has been widely adopted to characterise climatic, hydrologic and other environmental cues in ecology (Firman, Rubenstein, Moran, Rowe, & Buzatto, 2020; Wingfield et al., 1993). We calculated predictability as per Jiang et al. (2017), creating frequency tables for temperature and precipitation events using monthly time steps and set bins for climatic variables. Decisions around the binning of continuous climatic variables are fundamental to this method of calculating predictability (Jiang et al., 2017). Given temperature predictability tends to vary fairly consistently along a latitudinal gradient globally (Jiang et al., 2017), we chose to bin temperature by fixed bins of 5°C with two bins of 10°C at each end of the scale to capture rare extreme values, resulting in a total of ten bins for temperature (i.e. breakpoints at −10, 0, 5, 10, 15, 20, 25, 30, 35, 40, 50). We binned precipitation with a base 3 exponential binning scheme, considering the large range of precipitation data and creating seven bins in total (0, 3^1, 3^2…3^7).

### 2.4 Data analysis

All analyses were performed in R version 4.0.4 (R Core Team, 2021). All data and analysis code are available at https://doi.org/10.5281/zenodo.5553530.

Trait-environment relationships were analysed according to the Community Weighted Means (CWM) approach detailed by ter Braak et al. (2018). AusPlots species cover-abundance scores were used to generate CWMs of flowering period lengths for each plot, which were then regressed using ordinary least squares (OLS) regression against temperature and precipitation means and predictability for those plots. MAP was log transformed (base 10) prior to analysis. To ensure that trait-environment relationships were robust, species cover-abundance scores were also used to calculate weighted Species Niche Centroids (SNC) for each species and each environmental variable, and these were regressed against species’ flowering period lengths. The highest p-value for each trait-environment relationship (CWM~enviro, SNC~trait) was retained (pmax) to screen for potential false positive relationships (ter Braak et al., 2018). To assess their combined predictive power we regressed significant climatic predictors against flowering period length CWMs using OLS multiple regression.

To further explore flowering period patterns, we compared flowering period lengths among biomes. Given the unequal numbers of AusPlots in different biomes, differences in CWM flowering period lengths between biomes were assessed using a Welch’s ANOVA for unequal variances with Games-Howell posthoc tests. We also compared the difference in flowering period lengths between woody and herbaceous species using Welch’s T-tests, with one t-test for all species pooled (n = 2790) and multiple t-tests with Bonferroni correction (alpha = 0.05/6 = 0.008) for species by biome (n = 87-1160). Data on woodiness were sourced from AusTraits (Falster et al., 2021). We also confirmed that species range size was positively correlated with flowering period length using OLS regressions for all available species (n = 2819) as an indication of the potential intraspecific variation in flowering phenology captured by species-level data. Range size data (as extent of occurrence, or EOO) was sourced from Gallagher et al. (2021).

## 3 RESULTS

### 3.1 Trait-environment relationships

Community weighted mean (hereafter ‘community’) flowering period lengths increased with MAT and decreased with MAP, precipitation predictability and temperature predictability, though no single relationship explained greater than 20% of community variation (Table 1, Figure 2). The relationship between community flowering period length and environmental variables was strongest for temperature predictability (R^2^ = 0.17, pmax < 0.001). MAT and MAP both explained just over 10% of variation in community flowering period lengths (R^2^ = 0.11, pmax < 0.001). The relationship between precipitation predictability and community flowering period lengths was weaker (R^2^ = 0.09 pmax < 0.001). All climatic predictors combined explained 29% of variation in community flowering period length (multiple linear regression *F*_4,572_ = 59.53, p < 0.001, R^2^ = 0.29). All climatic predictors contributed significantly to the multiple regression (p < 0.04 in each case), with no multicollinearity among predictors (VIF < 2.2 in each case, correlations −0.11 – 0.71; Figure S3).

**Figure 2.**
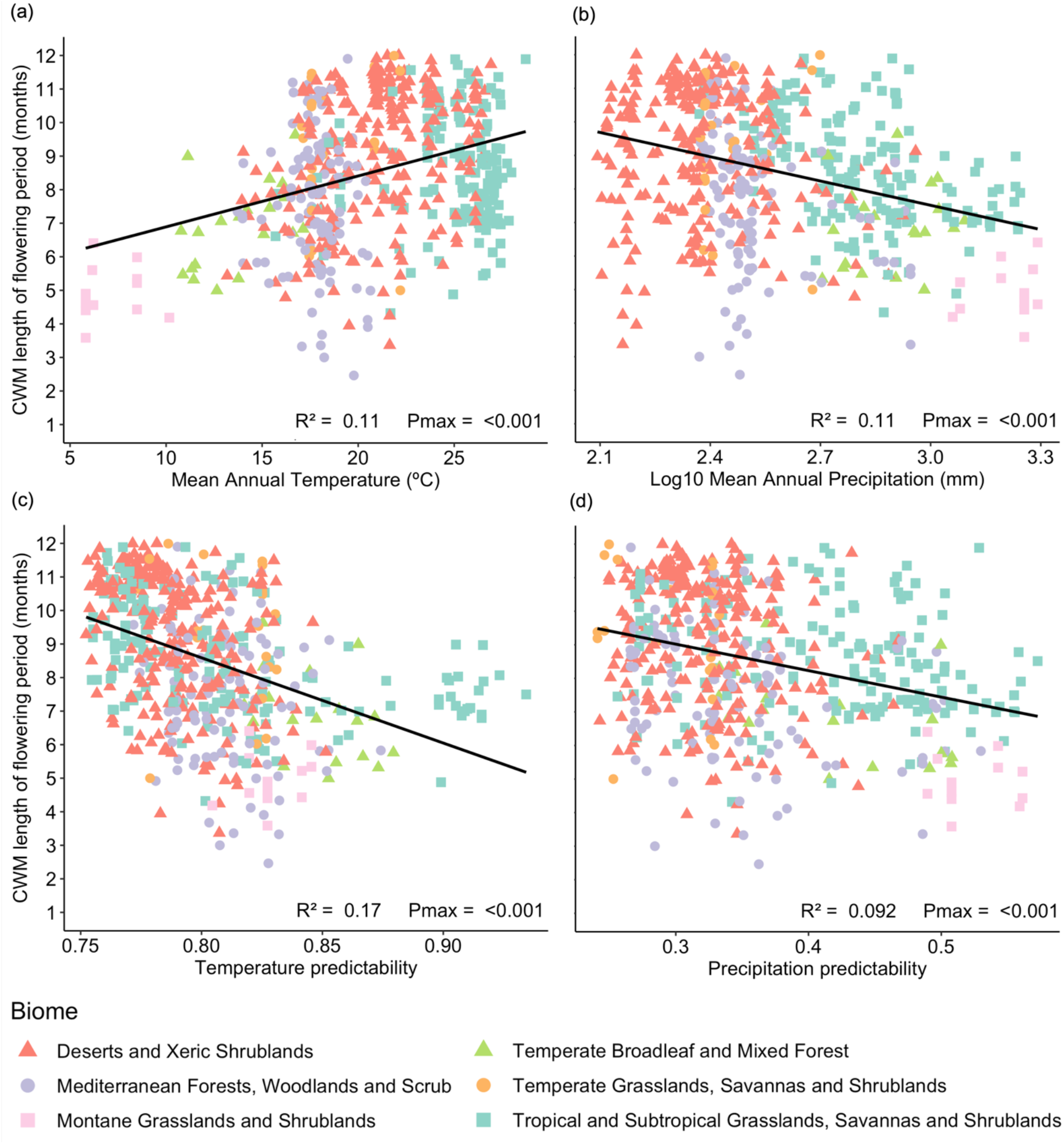
Relationships between mean annual temperature (°C) (a) mean annual precipitation (mm) (b), temperature predictability (c), precipitation predictability (d), and community weighted means (CWM) of the length of flowering periods (months). Pmax values report the highest P value for both SNC and CWM regressions.

**Table 1.**
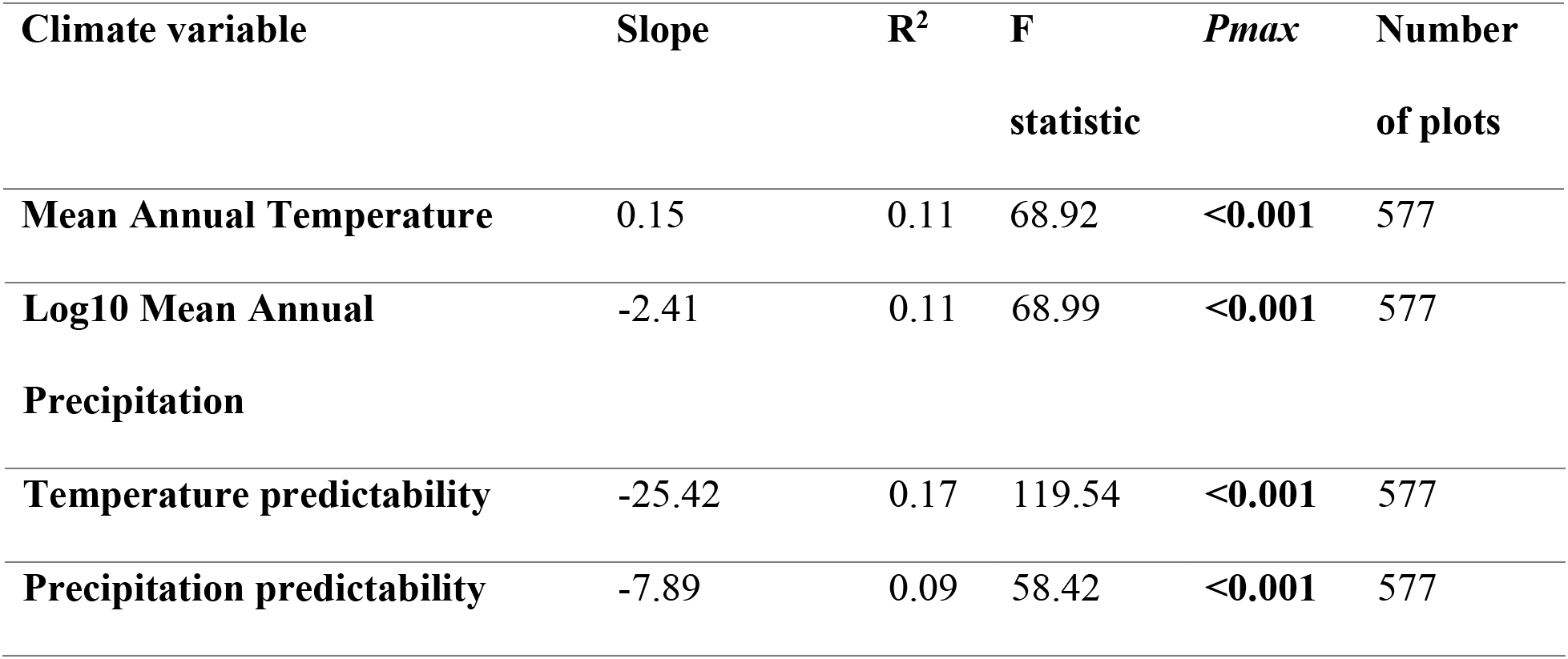
Results from ordinary least squares regressions of community weighted mean flowering period length versus climatic variables. Pmax reports the highest p-value from CWM and SNC regressions for the same climate variable.

### 3.2 Flowering periods by biome

Community flowering period lengths differed significantly by biome (Welch’s ANOVA for unequal variances F_5,64.82_ = 63.25, *P* < 0.001; Figure 3). Community flowering periods were longest on average in Deserts and Xeric Shrublands, closely followed by Temperate Grasslands, Savannas and Shrublands and Tropical and Subtropical Grasslands, Savannas and Shrublands (Figure 3; Table S2). Community flowering periods were shortest in Montane Grasslands and Shrublands (Figure 3; Table S2).

**Figure 3.**
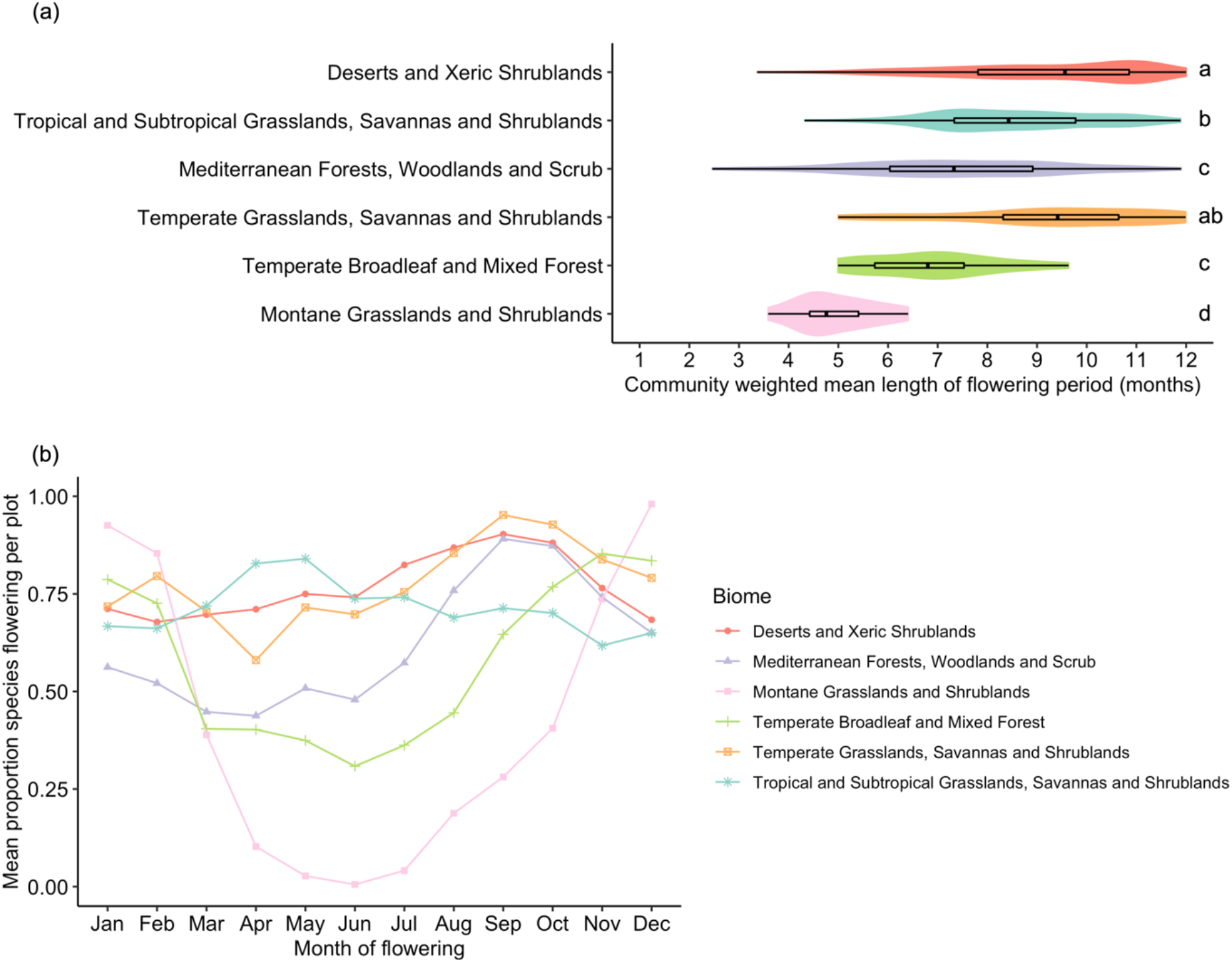
Flowering periods by biome: a) community weighted mean flowering period lengths. Letters indicate significantly different groups according to Games-Howell posthoc tests; b) monthly pattern of flowering as mean proportion of species cover flowering per site per month.

Montane Grassland and Shrubland sites show a strongly seasonal pattern of flowering, followed by Temperate Broadleaf and Mixed Forest (Figure 3). In contrast, Tropical and Subtropical Grasslands, Savannas and Shrublands; Temperate Grasslands, Savannas and Shrublands; and Desert and Xeric Shrubland biomes all show aseasonal patterns of flowering (Figure 3). When considering the geographic distribution of community flowering period lengths, central and northern Australia show generally longer community flowering periods, with shorter community flowering periods in southwest Western Australia and south-eastern Australia (Figure S5).

Flowering period lengths across all species and among biomes are shown in Figure 4. The plant families contributing most species, occurrences and proportionate cover in study plots were Fabaceae (414 species, 1626 occurrences, 88 cumulative proportional cover), Poaceae (374 species, 2743 occurrences, 196 cover) and Myrtaceae (287 species, 1033 occurrences, 125 cover; Table S1, Figure 4). Some families had relatively low species richness but high cover, including Casuarinaceae (13 species with 15 cover) and Scrophulariaceae (54 species with 11 cover). The distribution of flowering period lengths has peaks at three months, six months and twelve months, with most values falling between three and six months (Figure 4). Flowering periods of twelve months were particularly common for Poaceae and Chenopodiaceae species, whilst flowering periods of three to six months were more common for species in the Fabaceae and Myrtaceae (Figure 4). Different species flowering period lengths among AusPlots biomes therefore reflect the uneven distribution of plant families among biomes (Figure 4).

**Figure 4.**
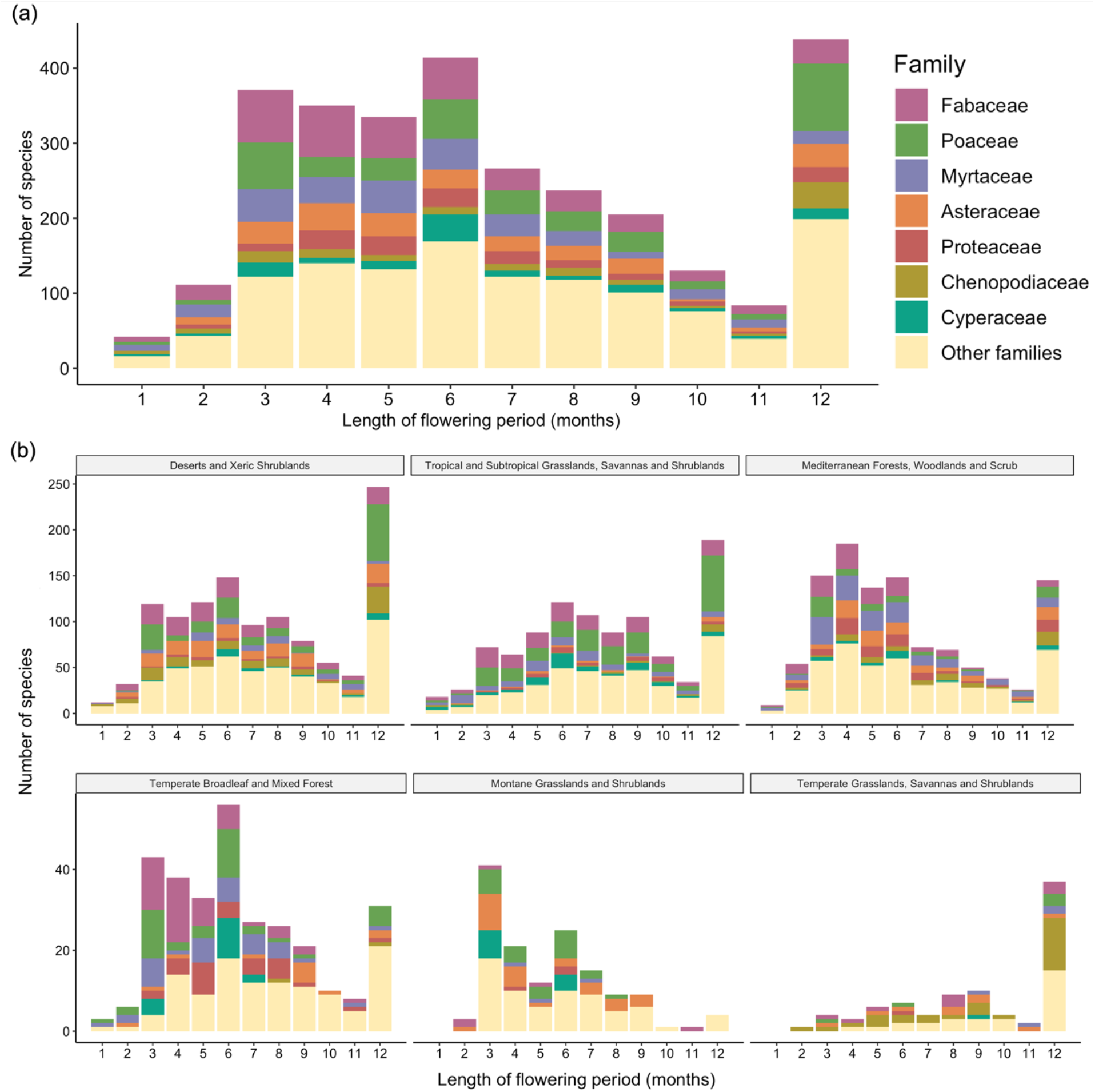
The distribution of species flowering period lengths, coloured by family: a) total and b) separated by biome. Note in (b) the different scales between the larger (top row) and smaller (bottom row) biomes.

Flowering periods were longer in species with larger extents of occurrence (R^2^ = 0.2, p < 0.001; Figure 5). Mean flowering periods were longer for herbaceous species (mean = 6.8) than woody species (mean = 6.5; t_2786_ = −2.73, p = 0.01; Figure S4). Mean flowering periods did not differ significantly between woody and herbaceous species within different biomes (alpha with Bonferroni correction = 0.008, p = 0.03-0.49).

**Figure 5.**
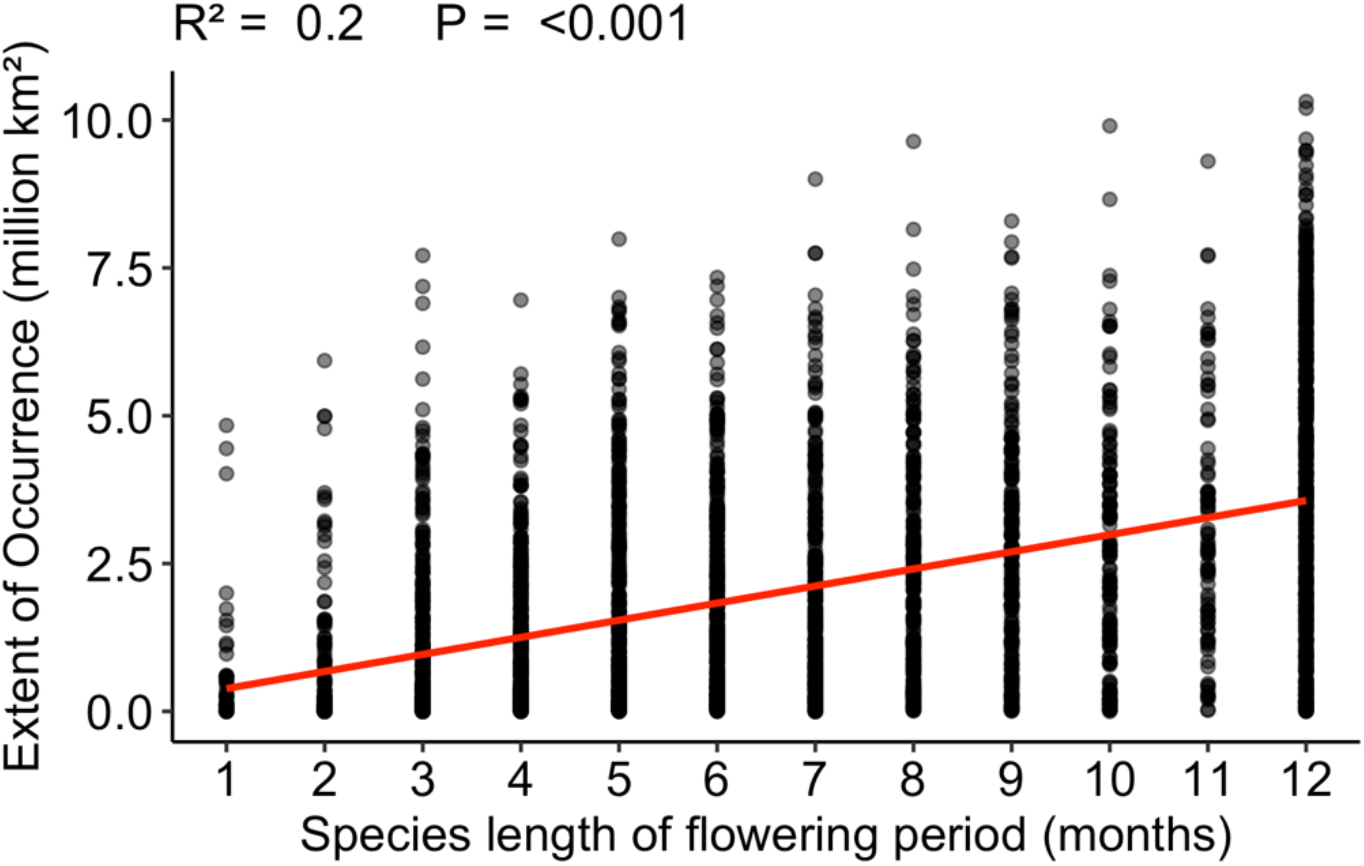
Species length of flowering period (months) against species extent of occurrence (million km^2^).

## 4 DISCUSSION

We show that climate plays a significant role in determining flowering period of plant communities, not just their constituent species, across six biomes, 23°C of MAT and 1,800 mm of MAP variation. Biome level differences in community flowering periods are driven in part by temperature and precipitation, both means and predictability. Four climate variables explained 29% of variation in community flowering period lengths (i.e. MAT, MAP, and the predictability of temperature and precipitation). As hypothesised, plant communities with higher MAT and lower MAP typically exhibit longer mean flowering periods, whereas plant communities with predictable temperatures and precipitation exhibit shorter mean flowering periods. While the relationship with temperature predictability was the strongest observed, the relationship with precipitation predictability was weaker than those with climatic means, perhaps due to the extreme variability and low predictability observed in precipitation across the Australian continent (Table 1). Our results show that shifts in flowering period with climate previously documented at the species level also operate in plant communities with implications for community assembly processes under both current and future climates.

Community flowering responses to climate are a product of the flowering phenologies of constituent species, which in turn depend on the flowering phenologies of constituent populations and individuals (Craine et al., 2012; Primack, 1985). Localised climatic conditions directly shape the flowering periods of the plant populations in an area, contributing to intraspecific variation, which then affects the flowering period recorded at the species level (Craine et al., 2012; Park, 2014). Though we could not test it directly, the effect of intraspecific variation on flowering period length is suggested in our results, as species with larger ranges have longer flowering periods (Figure 5). This illustrates how the use of species-level flowering periods as opposed to site-specific data may shape our results: as species range size increases, the specificity of flowering time observations decreases. Larger species’ ranges encompass a broader array of climatic conditions, which should lead to longer periods of time in which different populations may experience suitable conditions for flowering. Thus, intraspecific responses to climate likely affect our results indirectly, shaping the species flowering periods that in turn contribute to community level flowering periods.

At the community scale, climate conditions can influence the co-occurrence of species with particular flowering periods via environmental filtering (Du et al., 2020; Park, 2014). Though the influence of climate is typically weaker when examining interspecific relative to intraspecific flowering times, composition-derived shifts in flowering time can explain up to 49.3% of community phenological variation (Park, 2014). Phenology can be a major determinant of species distributions, setting geographic limits on the environmental conditions a species requires to complete its life cycle (Chuine, 2010). In our study different plant families predominate in different biomes, and these compositional shifts correspond with shifts in flowering periods among biomes (Figure 3, Figure 4). In addition, community mean flowering periods vary with climate, suggesting that flowering phenology may be one of several traits determining species co-occurrence in plant communities, along with more commonly investigated traits such as plant height and specific leaf area (Guerin et al., In review). This is supported by Du et al. (2020)’s finding that flowering and fruiting phenology varies with environment across China, and shows that climate-community phenology relationships can be detected even in local, co-occurring plant communities, despite the influence of stochastic events on local community assembly (Bruelheide et al., 2018). As such, our results clearly demonstrate the signal of environmental filtering in community flowering phenology, as different flowering strategies predominate across the breadth of plant communities and biomes explored.

### 4.2 Flowering period as a “response” trait

Flowering periods are longest in Desert and Xeric Shrubland communities, and in communities with low and unpredictable MAP. This reflects longstanding observations about the flowering phenology of desert communities, which is typically opportunistic in response to sporadic rainfall (Noy-Meir, 1973). The long flowering periods of desert biomes do not imply long flowering durations. Instead, longer flowering periods reflect the fact that desert species flower at any time of year in response to rainfall, which shows high inter-annual variability across Australia’s arid regions (Friedel et al., 1993; King et al., 2014). For plants to be able to meet the physiological costs of flower production and maintenance (both water and carbon, see Roddy *et al*. (2020)), and resulting seed production, they must respond to water when it is available. Plants respond to this unpredictable rainfall differently: desert annuals and herbaceous perennials often germinate, flower and fruit following rainfall, with annuals completing their full life cycle while soil moisture is available (Nano & Pavey, 2013; Noy-Meir, 1973). Woody species typically have deeper root systems with access to more stable soil moisture, and can thus access resources to flower in more predictable windows, but still respond to stochastic rainfall events for flowering and reproduction (Friedel et al., 1993, 1994; Nano & Pavey, 2013; Noy-Meir, 1973). These differences in woody and herbaceous species’ flowering may explain the slightly longer flowering periods found for herbaceous species, which showed a larger proportion of species with 12 month flowering periods than woody species (Figure S4), though this relationship did not hold within Deserts and Xeric Shrublands or any other biome.

In contrast to desert communities, mean flowering periods are shorter in Montane Grasslands and Shrublands, and in communities with low MAT, high MAP and predictable temperature and precipitation. Alpine plant communities experience strong climatic boundaries, with low temperatures and snow cover in the winter months preventing plant growth or reproduction. These strong climatic boundaries limit the window for flowering, pollination and seed production in alpine plant communities, which must be completed before autumn snowfall (Inouye & Pyke, 1988). Reflecting this, alpine plant communities experience the most seasonal flowering of any Australian biome, with peak flowering in December-January and no flowering in June, the month of the Southern Hemisphere’s winter solstice (Figure 3). The strength and specificity of this flowering pattern also reflects the smaller ranges of Australian alpine species (R. V. Gallagher, 2016). Australia’s montane biome covers a small proportion of the country’s terrestrial surface area (~0.16%, Figure 1) and is a centre of floral endemism in Australia (Crisp, Laffan, Linder, & Monro, 2001). Our findings confirm the combination of highly seasonal flowering, tight climate relationships and high rates of endemism which have made montane biomes the subject of intense research into the impacts of climate change on flowering phenology in recent decades (CaraDonna et al., 2014; R. Gallagher, Hughes, & Leishman, 2009). Some impacts of climate change on flowering phenology in Australian montane habitats have been detected, and these may lengthen community flowering periods in this biome in the future (R. Gallagher et al., 2009; Green, 2010).

Community mean flowering periods decreased with increasing predictability of both temperature and precipitation, as hypothesised. Precipitation predictability had less explanatory power than climatic means, while temperature predictability explained the most variance in community flowering periods (Table 1). Flowering is highly responsive to temperature cues, with flowering in many species initiated by increases in ambient temperatures (Capovilla, Schmid, & Posé, 2015). It is thus unsurprising that more predictable temperature cues equal more regular, and thus shorter, community flowering periods, although Australian temperatures are highly predictable compared to other regions of the world (Jiang et al., 2017). In contrast, precipitation in Australia is highly variable both geographically and year-to-year, driven by climatic modes such as the El Niño-Southern Oscillation, and this contributes to low levels of precipitation predictability (King et al., 2014). Temperate Broadleaf and Mixed Forest biomes in Australia, for example, are globally unique for their low precipitation predictability, and in particular their low precipitation contingency (Jiang et al., 2017). Australian vegetation is correspondingly opportunistic, with growth and flowering events often closely tracking water availability (Duursma et al., 2016; Nano & Pavey, 2013). Though community flowering periods decrease with precipitation predictability as predicted, this relationship was weaker than that with other climatic predictors, perhaps due to the extreme heterogeneity of precipitation across the Australian continent. Overall the relationship between climatic predictability and community plant phenology across Australia suggests climatic factors shaping plant community assembly beyond the climatic means typically considered.

### 4.3 Flowering period as an “effect” trait

What do our results about flowering period imply for pollinators and pollination? Pollination is spatially heterogeneous: for example, wind pollination is thought to be more common in areas with lower MAT and MAP (Rech et al., 2016). For animal-pollinated species, different pollinator assemblages are active in different areas and different climatic conditions (Ollerton, 2017). Areas with higher MAT likely have more months of the year in which pollinator species are active (Primack & Inouye, 1993), and thus increased flowering periods in these communities is likely matched by increased windows of pollinator activity.

Relationships between pollinator activity and precipitation are more complicated. Though areas with higher precipitation have increased water availability which can increase floral traits associated with pollinator attraction and reward, rainfall itself typically impedes pollinator activity, diluting flower nectar, degrading pollen and preventing insect pollinators from flying (Lawson & Rands, 2019). Pollinator activity likely varies with climatic predictability much as flowering periods do, though pollinator phenology is less frequently or consistently studied (Neave, Brown, Batley, Rao, & Cunningham, 2020). In desert biomes, for example, bird abundance and species richness tracks unpredictable rainfall (Jordan, James, Moore, & Franklin, 2017), and pollinators in cold or montane environments experience similar periods of reduced activity, either migrating away or else overwintering as larvae during the cold months (Inouye & Pyke, 1988; Stemkovski et al., 2020). Thus, climate shapes community flowering periods but also the activity of pollinators that visit flowers, not to mention the activity of the many florivorous animals that do not effect pollination (e.g. see McCall & Irwin, 2006).

### 4.4 Implications and future directions

Community flowering strategies may shift with climate change, either as species adapt to new conditions or as community composition changes via localised extinctions and range shifts. In Australia climate change is causing higher temperatures overall, with an increase in heavy precipitation in northern Australia and an increase in drought in southern Australia (IPCC, 2021). Communities with shorter flowering periods will be more susceptible to the impacts of current and future climate change, as mismatches in the timing of flowering, pollinator emergence and climatic conditions over time may select for communities with longer, more responsive flowering periods (e.g. Stemkovski *et al*., 2020). Indeed, there are already reports that lower and less predictable rainfall is affecting plant community composition through dieback in southwest Australia (Hoffmann et al., 2019), and that higher temperatures are shifting flowering dates in alpine southeast Australia (R. Gallagher et al., 2009; Hoffmann et al., 2019).

Flowering is just one part of a plant’s reproductive phenology, and flowering phenology is just one aspect of a plant’s floral strategy. Seed size may influence the timing of fruiting and flowering, as flowers must be pollinated in time to allow suitable conditions for fruit development, which takes longer in larger-seeded species, and seed dispersal (Chuine, 2010; Du et al., 2020). Evidence for this hypothesis is equivocal, however, and recent field investigations in montane habitats found no association between phenological events and seed size, though they did find a strong association with plant height (Liu et al., 2021). A landscape scale comparison between plant traits, fruiting time and flowering period would require either more specific measures of population flowering duration, or else measurement only in strongly seasonal environments where flowering periods experience definite constraints. A more fruitful approach in aseasonal landscapes might be to investigate community-level variation in other floral traits, in particular traits related to trade-off spectra such as floral longevity, floral mass or floral mass per area (Roddy et al., 2020).

## 5 Conclusion

Climate has long been known to affect plant strategies across biomes. Here we have shown that climate similarly contributes to strategies around the timing of plant flowering. Plant communities in climatically predictable areas, with higher mean precipitation and lower mean temperatures, favour shorter, more concentrated flowering periods. Species in these communities likely time their flowering to match pollinator activity and optimal conditions for pollination and seed development. In contrast, plant communities in areas with unpredictable climates, with lower mean precipitation and higher mean temperatures, have longer, more dispersed flowering periods, as species in these harsher conditions must respond whenever water is available to enable flowering. Filtering for these divergent flowering strategies may limit which species can co-exist in communities, resulting in signals of flowering in the processes of community assembly. Future studies may further reveal how different flowering strategies affect pollination, plant reproduction and community turnover, as well as the availability of floral resources across the landscape.

## Acknowledgements

R.E.S. was supported by funding through the Australian Government’s Research Training Program. MJ acknowledges the Australian Research Council DECRA fellowship (DE210101654). We thank all contributors to the AusTraits plant trait database, particularly taxonomists and their supporting institutions for their long-term work describing the flora of Australia. We also acknowledge all contributors to TERN AusPlots for their work in creating an excellent source of reliable plant abundance data for macroecological work in Australia.

## Data availability statement

All data used in this study and primary analysis R code are available via an archived GitHub repository at https://doi.org/10.5281/zenodo.5553530.

## Biosketch

Ruby E. Stephens is a plant ecologist with broad interest in the macroecology of plants and pollination. This work is part of her PhD on the macroecology and macroevolution of flowers and floral traits across Australia and globally. She and other authors collaborate on studies of plant traits, macroecology and macroevolution via the AusTraits project (https://austraits.org/), the Terrestrial Ecosystem Research Network (https://www.tern.org.au/) and across their respective universities and lab groups.

## Author contributions

R.E.S., R.V.G. and H.S. conceived the project; G.R.G. provided example code and assistance with AusPlots data, D.F. assisted with taxonomic name matching, M.J. calculated climate data. R.E.S. analysed the data; R.E.S. led the writing with assistance and review from all other authors.

