## Supplementary Material for "Climate shapes flowering periods across plant communities"

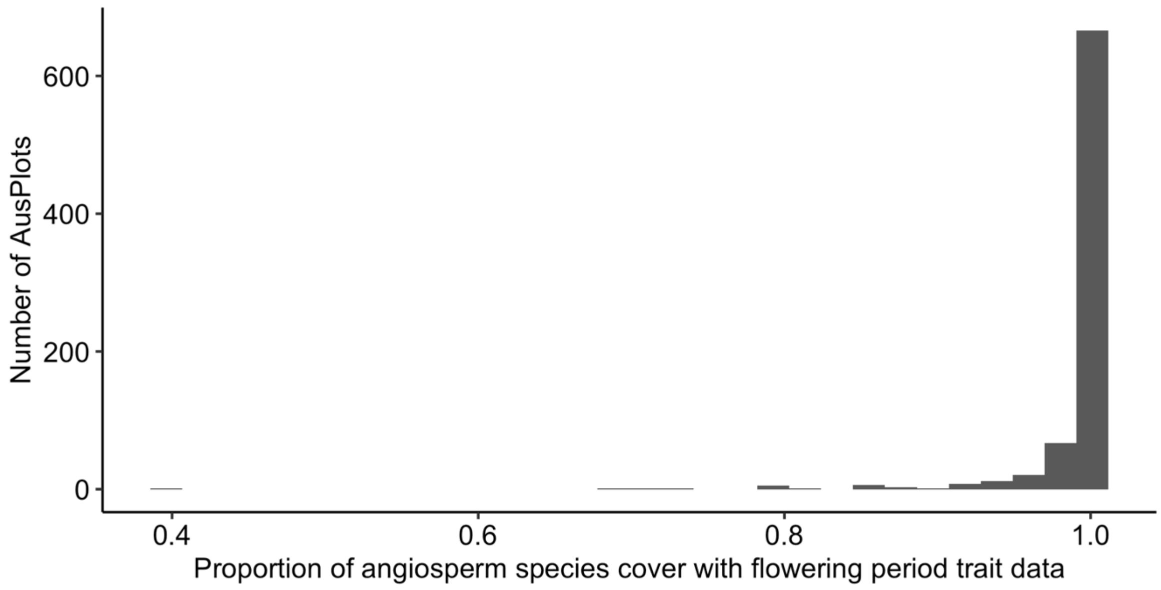


*Figure S1 The proportion of angiosperm species cover with matching flowering period trait data across available AusPlots. Only AusPlots with ≥ 80% of angiosperm species cover matched to trait data were included in the analysis.*

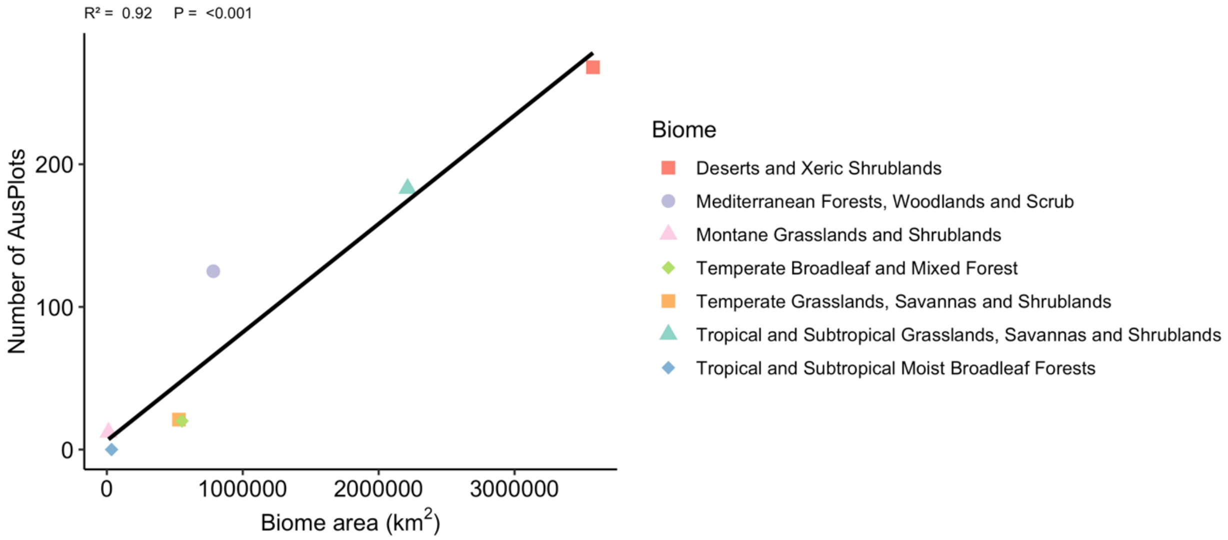


*Figure S2 The number of sites in each biome included in our analysis correlated with the terrestrial area in kilometres squared of each biome across Australia (ordinary least squares regression $R^2 = 0.92$, $P < 0.001$). Biomes are based on the Dinerstein et*

*al. 2017 global terrestrial ecoregions aligned to the Australian Interim Biogeographic Regionalisation for Australia v7 (Australian Department of the Environment and Energy, 2016).*

*Table S1 The top 10 plant families in study plots by number of species, with total number of occurrences and cumulative proportion of cover (i.e. the weight used in Community Weighted Means) summed across all plots.*

| **Family** | **Number of species** | **Number of occurrences** | **Proportion cover** |
|---|---|---|---|
| **Fabaceae** | 414 | 1626 | 88 |
| **Poaceae** | 374 | 2743 | 196 |
| **Myrtaceae** | 287 | 1033 | 125 |
| **Asteraceae** | 229 | 744 | 16 |
| **Proteaceae** | 154 | 461 | 9 |
| **Chenopodiaceae**[1] | 124 | 1294 | 69 |
| **Cyperaceae** | 124 | 333 | 9 |
| **Malvaceae** | 86 | 440 | 9 |
| **Goodeniaceae** | 64 | 181 | 3 |
| **Scrophulariaceae** | 54 | 272 | 11 |
| **Other families** | 1073 | 3702 | 94 |

*[1] The family Chenopodiaceae is still recognised by the Australian Plant Census despite being included in Amaranthaceae elsewhere.*

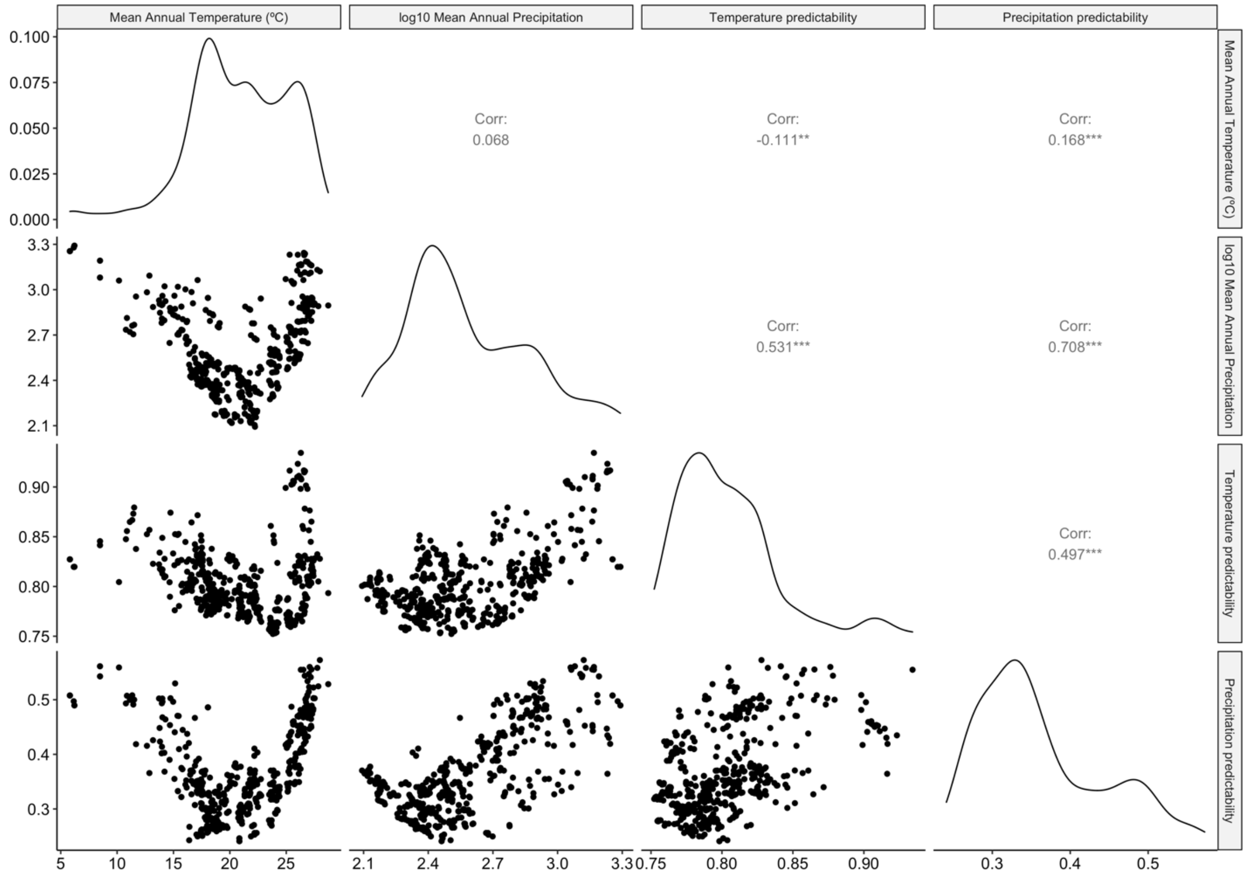


*Figure S3 Pairwise correlation plot for climatic predictors including Mean Annual Temperature (°C), log10 Mean Annual Precipitation, Temperature Predictability and Precipitation Predictability*

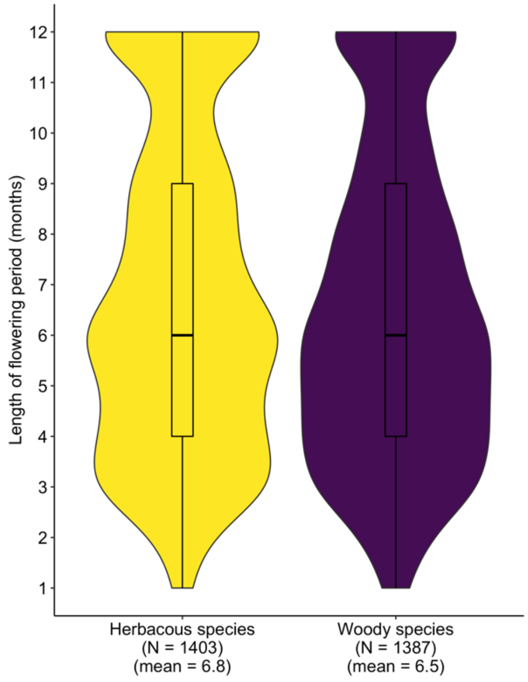


*Figure S4 Length of flowering period (months) for herbaceous and woody species.*

*Table S2 Results from Games-Howell posthoc test of differences in mean community flowering periods among biomes. Adjusted p-values <0.05 indicate that the mean community flowering period length differs significantly between the two given biomes.*

| **Biome 1** | **Biome 2** | **Mean difference** | **Adjusted p-value** |
|---|---|---|---|
| Deserts and Xeric Shrublands | Mediterranean Forests, Woodlands and Scrub | -1.77 | <0.001 |
| Deserts and Xeric Shrublands | Montane Grasslands and Shrublands | -4.28 | <0.001 |
| Deserts and Xeric Shrublands | Temperate Broadleaf and Mixed Forest | -2.30 | <0.001 |
| Deserts and Xeric Shrublands | Temperate Grasslands, Savannas and Shrublands | 0.12 | 1.000 |
| Deserts and Xeric Shrublands | Tropical and Subtropical Grasslands, Savannas and Shrublands | -0.65 | 0.001 |

| | | | |
|---|---|---|---|
| Mediterranean Forests, Woodlands and Scrub | Montane Grasslands and Shrublands | -2.51 | <0.001 |
| Mediterranean Forests, Woodlands and Scrub | Temperate Broadleaf and Mixed Forest | -0.53 | 0.612 |
| Mediterranean Forests, Woodlands and Scrub | Temperate Grasslands, Savannas and Shrublands | 1.89 | 0.004 |
| Mediterranean Forests, Woodlands and Scrub | Tropical and Subtropical Grasslands, Savannas and Shrublands | 1.12 | <0.001 |
| Montane Grasslands and Shrublands | Temperate Broadleaf and Mixed Forest | 1.98 | <0.001 |
| Montane Grasslands and Shrublands | Temperate Grasslands, Savannas and Shrublands | 4.39 | <0.001 |
| Montane Grasslands and Shrublands | Tropical and Subtropical Grasslands, Savannas and Shrublands | 3.63 | <0.001 |
| Temperate Broadleaf and Mixed Forest | Temperate Grasslands, Savannas and Shrublands | 2.42 | 0.001 |
| Temperate Broadleaf and Mixed Forest | Tropical and Subtropical Grasslands, Savannas and Shrublands | 1.66 | <0.001 |
| Temperate Grasslands, Savannas and Shrublands | Tropical and Subtropical Grasslands, Savannas and Shrublands | -0.76 | 0.536 |

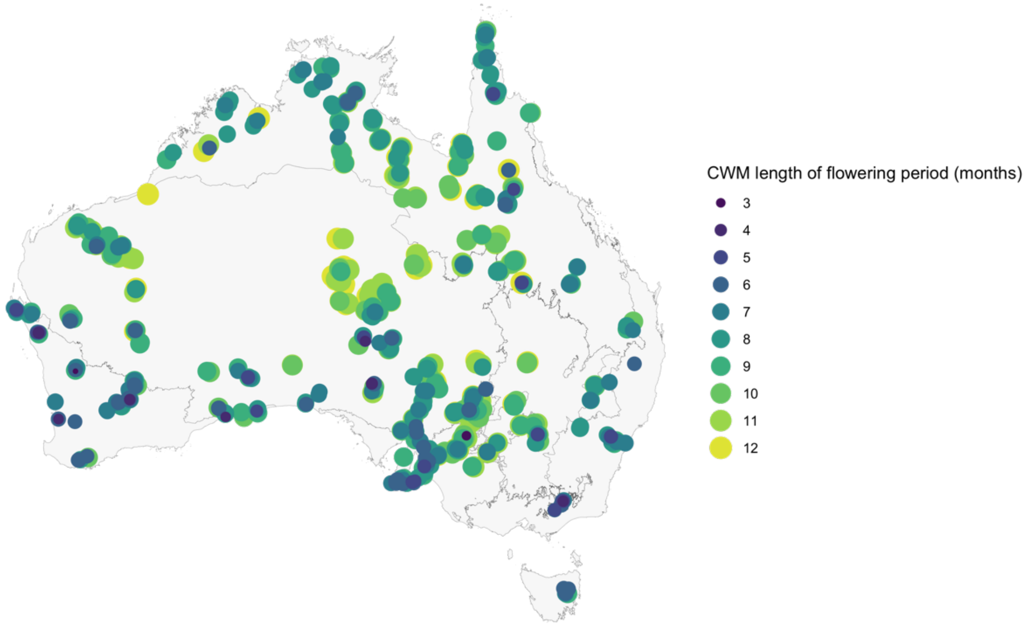


*Figure S5 Map showing the Community Weighted Mean (CWM) length of flowering period (months) for 629 AusPlots across Australia.*
